# A precision overview of genomic resistance screening in isolates of *Mycobacterium tuberculosis* using web-based bioinformatics tools

**DOI:** 10.1101/2023.01.10.523521

**Authors:** Gabriel Morey-León, Paulina M. Mejía-Ponce, Juan Carlos Granda Pardo, Karen Muñoz-Mawyin, Juan Carlos Fernández-Cadena, Evelyn García-Moreira, Derly Andrade-Molina, Cuauhtémoc Licona-Cassani, Luisa Berná

**Author notes:** Corresponding author. Luisa Berná. These authors contributed equally to this work (Gabriel Morey-León, Paulina M. Mejía-Ponce).

## Abstract

Tuberculosis (TB) is among the most deadly diseases that affect worldwide, its impact is mainly due to the continuous emergence of resistant isolates during treatment due to the laborious process of resistance diagnosis, non-adherence to treatment and circulation of previously resistant isolates of *Mycobacterium tuberculosis*. The aim in this study was evaluate the performance and functionalities of web-based tools: Mykrobe, TB-profiler, PhyReSse, KvarQ, and SAM-TB for detecting resistance in isolate of *Mycobacterium tuberculosis* in comparison with conventional drug susceptibility tests. We used 88 *M. tuberculosis* isolates which were drug susceptibility tested and subsequently fully sequenced and web-based tools analysed. Statistical analysis was performed to determine the correlation between genomic and phenotypic analysis. Our data show that the main sub-lineage was LAM (44.3%) followed by X-type (23.0%) within isolates evaluated. Mykrobe has a higher correlation with DST (98% of agreement and 0.941Cohen’s Kappa) for global resistance detection, but SAM-TB, PhyReSse and Mykrobe had a better correlation with DST for first-line drug analysis individually. We have identified that 50% of mutations characterised by all web-based tools were canonical in *rpoB, katG, embB, pncA, gyrA* and *rrs* regions. Our findings suggest that SAM-TB, PhyReSse and Mykrobe were the web-based tools more efficient to determine canonical resistance-related mutations, however more analysis should be performed to improve second-line detection. The improvement of surveillance programs for the TB isolates applying WGS tools against first line drugs, MDR-TB and XDR-TB are priorities to discern the molecular epidemiology of this disease in the country.

**Importance:** Tuberculosis, an infectious disease caused by *Mycobacterium tuberculosis*, which most commonly affects the lungs and is often spread through the air when infected people cough, sneeze, or spit. However, despite the existence of effective drug treatment, the patient adherence, long duration of treatment, and late diagnosis, have reduced the effectiveness of therapy and raised the drug resistance. The increase in resistant cases, added to the impact of the COVID-19 pandemic, have highlighted the importance of implementing efficient and timely diagnostic methodologies worldwide. The significance of our research is in evaluating and identifying the more efficient and friendly web-based tool to characterise the resistance in *Mycobacterium tuberculosis* by whole genome sequencing, which will allow apply it more routinely to improve TB strain surveillance programs locally.

## Introduction

Tuberculosis (TB), caused by *Mycobacterium tuberculosis* (Mtb), is one of the top 10 causes of death worldwide, in 2021 the World Health Organization estimated 10.6 million people were infected and 1.6 million people died (1). Despite many innovations in tuberculosis in diagnosis, drug resistance, drug therapy, prevention and control programs, several challenges, such as patient adherence, long duration of treatment, and late diagnosis, have reduced the effectiveness of TB therapy (2). The increase in resistant TB patients is not only due to exposure to multidrug-resistant (MDR) and extensively resistant strains (XDR), but also due to late or inadequate diagnosis, ineffective treatment or poor adherence to treatment (2). The characteristic slow growth of Mtb is a constraint to the rapid diagnosis of anti-TB drug resistance and aggravates the situation by increasing the incidence of MDR and XDR-TB in the world. On the other side, the accumulation of point mutations in coding regions for drug targets and/or drug-converting enzymes is a major mechanism for acquiring resistance in MTB (3). Therefore, providing adequate treatment based on the rapid detection of resistant strains, as well as the identification of transmission clusters are crucial to effective treatment and prevention of onward transmission (4), for which the use of procedures that reduce the time of diagnosis is recommended. Traditional TB drug susceptibility testing (DST) relies on solid or liquid culture, which may take a minimum of 14 days or even more than a month to yield results. On the contrary, whole-genome sequencing (WGS) technologies can determine drug susceptibility in about 8–9 days (5).

Inside all diagnostic innovations in TB for resistance detection, it has been shown that WGS as a molecular diagnostic tool can be used to detect/rule out resistance to all drugs simultaneously (6–8). It also allows lineage identification, transmission tracing and outbreak definition (7–11) and is a faster and more effective method than traditional phenotypic DST (12). The use of WGS as a diagnostic tool goes hand in hand with the development and availability of a wide range of bioinformatics tools that facilitate processing and obtaining results quickly and efficiently. These include web-based tools such as TBProfiler(13), KvarQ(14), TGS-TB(15), Mykrobe Predictor TB(16), CASTB(17), PhyTB(18), ReSeqTB-UVP(19), GenTB(20), PhyResSE(21), SAM-TB(22) and others that have been developed to genotyping and drug resistance identification. The choice to use web tools for TB lineage and resistance characterisation is often based on ease of use, agreement in classification (lineage or resistance-associated to single nucleotide polymorphisms (SNPs)), feasibility of uploading information for batch analysis, availability to carry out the analysis with limited programming knowledge, user-friendly handling and easiest to understand results by technician. Nevertheless, some the web-based tool had some disadvantages: its not giving information about the quality of sequencing in terms of coverage and depth, not being able to make a phylogenetic tree with generated information, depending on the web access and on-line analysis, install the pipeline in standalone machine or server, require a lot of bioinformatics resources, among others. Despite this, the synergy of these tools with the WHO updated catalogue of resistance-associated mutations in TB (23) has allowed an easy implementation of WGS in surveillance programs of several countries based on efficient, economical and reliably results (24–26), allowing them to be considered as “useful” to guide diagnosis and treatment of this disease.

The incidence of tuberculosis in many countries, including Ecuador, has been rising since 2015 (27), also with the complexity of the COVID-19 pandemic. Moreover, the frequent emergence of drug-resistant strains is a serious global threat and poses significant challenges to public health, especially in low- and middle-income countries. For this reason, it is important to apply strategies that allow timely evaluation of resistance-conferring mutations in *Mycobacterium tuberculosis*, especially in local settings, where this information could assist in the development and adoption of rapid diagnostic and consequently, in the control of MDR-TB. This study is aimed at assessing the performance and functionalities of web-based tools Mykrobe, TB-profiler, PhyReSse, KvarQ, and SAM-TB for detecting resistance in isolate of *Mycobacterium tuberculosis* in comparison with conventional drug susceptibility tests (DST).

## Results

Resistance to antituberculosis drugs was evaluated by DST for 88 *M. tuberculosis* isolates obtained from patients between 2019 to 2021. The four first-line drugs (Rifampicin, Isoniazid, Pyrazinamide, and Ethambutol), three the second-line drugs (Streptomycin, and the fluoroquinolones: Levofloxacin and Moxifloxacin) and three injectable-drugs (Kanamycin, Amikacin or Capreomycin) were used to determine resistance. Of the total number of samples 22 isolates were identified as sensitive and 66 isolates were determined to be drug-resistant (DR-TB) and further classified as: MDR (n=46); Poly-resistant (n=13); resistant to RIF and fluoroquinolone (n=5); and resistant to RIF alone (n=2). Among the 88 samples, 26.1% were female, with a median age of 39.2 ± 14.5 years (range 21 -73 years) and 73.9% were male, with a median age of 40.3 ± 15.0 years (range 11 -74 years). Regarding the treatment scheme received, 37.5% of isolates come from treatment-naive cases (45.5% were resistant), 23.9% were in treatment (90.5% were resistant), and 38.6% come from previous-treatment cases (94.1% were resistant). Most resistant isolates (59.1%) were detected in Guayaquil, one of the most important economic city in Ecuador, followed by Babahoyo, El Empalme and Quito (4.5%, 3.4%, 2.3%, respectively). Only a few cases were detected in Chone, Duran, Guaranda, Machala, Nueva Loja (1.1% each) (S1 Table).

The phylogenetic relationships of the 88 isolates were evaluated from 5586 SNPs using the MTBseq pipeline. It was determined that there is a great diversity of circulating isolates in Ecuador, with LAM sublineages being the most representative (44.3%) followed by type X (23.9%) and type S (11.4%) Fig 1. There is no evidence of an association between sublineages and sensitivity to the drugs tested, but it is noteworthy that of the 11 S -type strains, 10 (88.9%) show MDR resistance and only one is sensitive. Based on the cluster analysis we identified 54 isolates (54/88, 61.4%) were grouped in 16 potentially molecular related transmission clusters (with at least two isolates, ≤12 SNP differences) and 34 singletons (S1 Table, Fig 1). In particular, seven clusters were considered small (< 3 isolates), eight medium (3 - 5 isolates) being the more prevalent the clusters 13 and 16 (five isolate each) and one largest cluster with 9 isolates (group 01). The XDR isolates were not defined in a single group and were distributed in the clusters 10 and 16 (two isolates in each). Similarly, the pre-XDR isolates were distributed in 4 clusters, two isolates in cluster 11 and one isolate in clusters 01, 08 and 10. The other pre-XDR isolates did not form clusters. The other three isolates were unique. Finally, the only large cluster, with nine isolates, consists entirely of the S-type sublineage.

**Figure 1.**
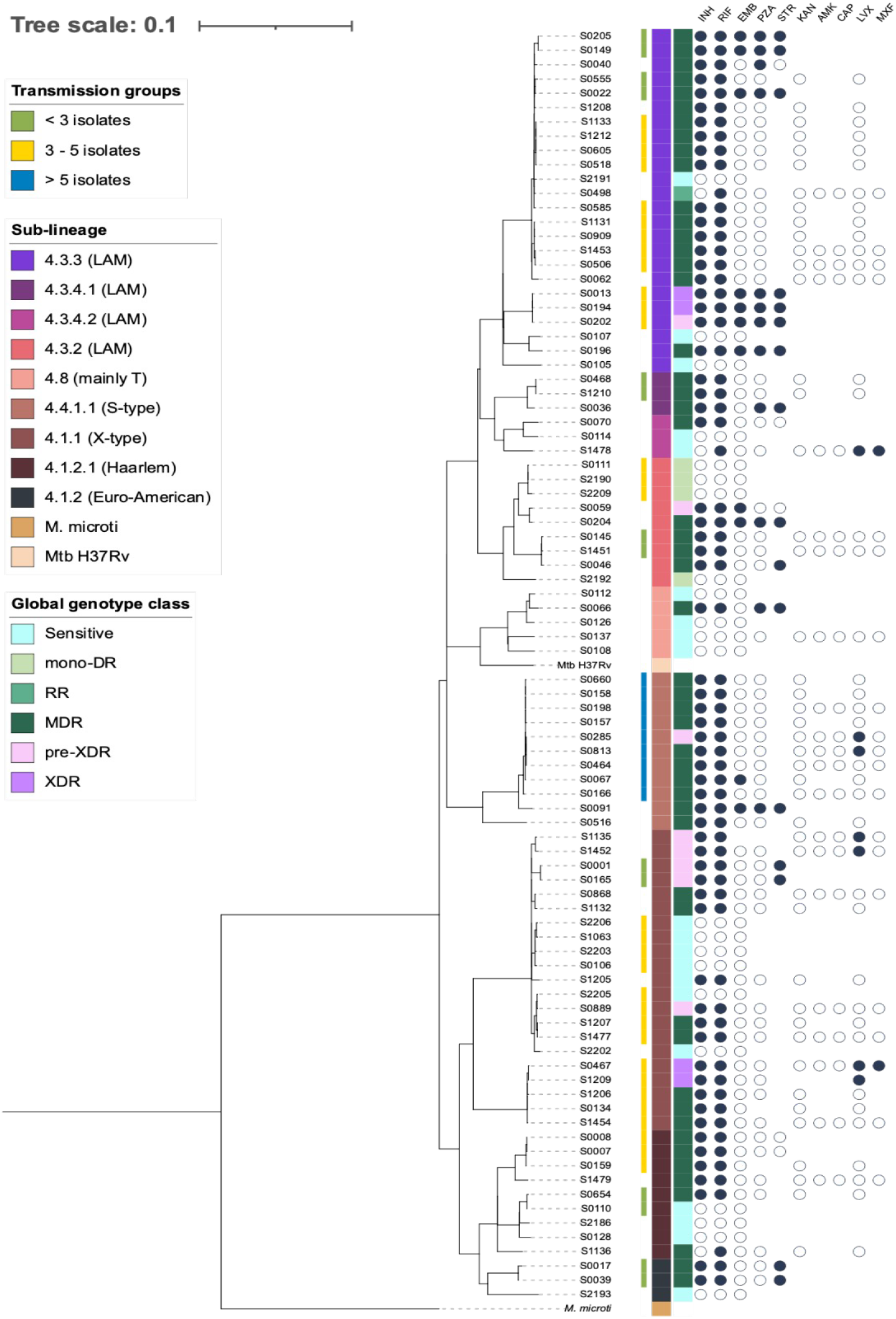
Phylogenetic reconstruction of the 88 Mtb isolates from Ecuador. A total of 5,586 SNPs were used to reconstruct the phylogenetic tree using the maximum likelihood method, the GTR substitution model and a bootstrap cutoff of 0.01. Metadata include i) transmission groups (lines green, yellow, and blue), ii) sublineage classification, iii) global genotypic class of drug resistance and iv) DST results of first- and second-line antibiotics (full circle = resistant; empty circle = sensitive; missing circle = not tested).

In order to evaluate the performance and functionalities of the web-based tools, the phenotypic resistance profile of 88 strains was compared with the characterised using the following web-based tools: TB-Profiler v2.8.13, PhyReSse v1.0, Mykrobe v0.10.0, KvarQ v0.12, and SAM-TB (S2 Table). Sensitivity, specificity, accuracy, percentage of agreement and Kappa coefficient (to measure inter-rater reliability) among other descriptive statistics are shown in Table 1. All the programs evaluated presented good sensitivity and specificity, although they showed different performance in the preconditions of resistance to specific drugs. We observed a highest level of agreement for isoniazid on all programs (agreement and Kappa Coefficient above 93% and k = 0,832); in the case of rifampicin, KvarQ was the lowest agreement (agreement 90.91% and k = 0,784). In the evaluation of the ability to detect specific resistances all tools show very good detection values for isoniazid and rifampicin (Table 2); whilst to ethambutol and pyrazinamide the detection was more variable, being adequate for Mykrobe, TB-profiler and SAM-TB but with lower quality values for KvarQ and PhyReSSe. In brief, TB-profiler and SAM-TB were the programs that showed the highest capacity to detect resistance to first-line antibiotics, while KvarQ was the lowest (Table 2).

**Table 1.** Comparative statistical analysis of five software for predicting anti-TB drug resistance.

| Statistic | Mykrobe | KvarQ | TB-Profiler | PhyReSse | SAM-TB |
| --- | --- | --- | --- | --- | --- |
| Sensitivity | 0.97 (0.89 to 0.99) | 0.92 (0.83 to 0.97) | 0.97 (0.89 to 1.00) | 0.97 (0.89 to 1.00) | 0.97 (0.89 to 1.00) |
| Specificity | 1.00 (0.85 to 1.00) | 0.86 (0.65 to 97) | 0.82 (0.60 to 0.95) | 0.82 (0.60 to 0.95) | 0.86 (0.65 to 97) |
| Positive Likelihood Ratio |  | 6.78 (2.36 to 19.44) | 5.33 (2.20 to 12.95) | 5.33 (2.20 to 12.95) | 7.11 (2.48 to 20.37) |
| Negative Likelihood Ratio | 0.03 (0.01 to 0.12) | 0.09 (0.04 to 0.21) | 0.04 (0.01 to 0.15) | 0.04 (0.01 to 0.15) | 0.04 (0.01 to 0.14) |
| Positive Predictive Value | 1.00 | 0.95 (0.88 to 0.98) | 0.94 (0.87 to 0.97) | 0.94 (0.87 to 0.97) | 0.96 (0.86 to 0.98) |
| Negative Predictive Value | 0.92 (0.74 to 0.98) | 0.79 (0.62 to 0.90) | 0.90 (0.69 to 0.97) | 0.90 (0.69 to 0.97) | 0.91 (0.71 to 0.97) |
| Accuracy (*) | 0.98 (0.92 to 1.00) | 0.91 (0.83 to 0.96) | 0.93 (0.86 to 0.97) | 0.93 (0.86 to 0.97) | 0.94 (0.87 to 0.98) |
| % of agreement | 0.98 | 0.91 | 0.93 | 0.93 | 0.94 |
| Cohen's kappa | 0.941 | 0.765 | 0.812 | 0.812 | 0.846 |
|  | Almost perfect agreement | Substantial agreement | Almost perfect agreement | Almost perfect agreement | Almost perfect agreement |

Numbers in brackets represent the corresponding 95% confidence intervals.

**Table 2.**
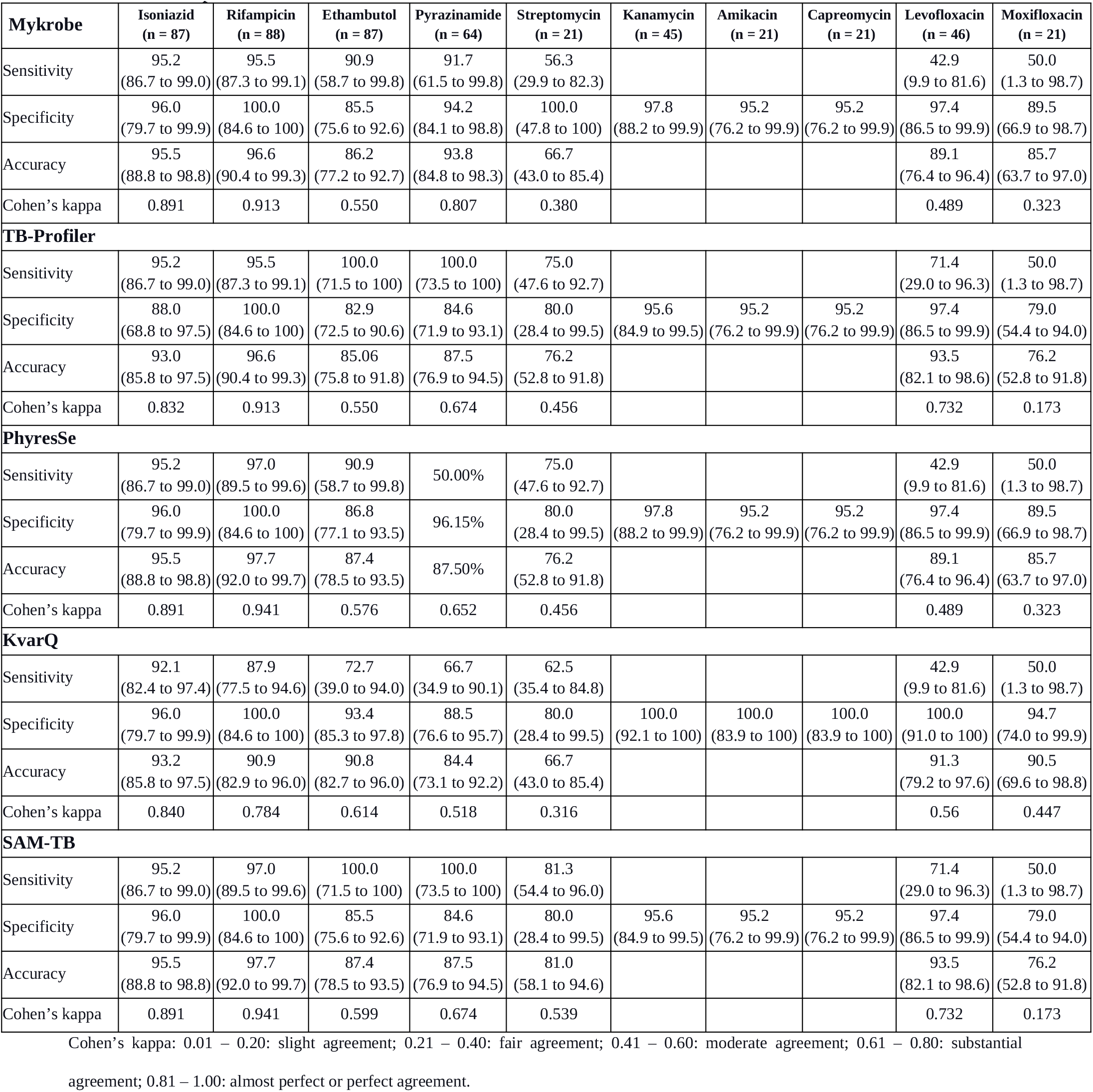
Sensitivity, specificity, and accuracy of ten drugs of anti-TB drug resistance analysed by WGS plus web-based tools compared with DST.

On the other hand, for streptomycin and moxifloxacin, the detection of resistance tends to a fair agreement with Mykrobe and KvarQ. All the tools analysed presented some problems in detecting resistance to fluoroquinolones of the second line, in particular to Moxifloxacin. Due only sensitive isolates were identified phenotypic in kanamycin, amikacin and capreomycin, some statistical parameter was not determined.

By genomic analysis of 88 phenotypic isolates, we identified genotypically 60.2% of strains MDR (53/88), 4.5% pre-XDR and XDR (4/88 each), 3.4% pre-MDR (3/88), and 19.6% Sensitive (19/88). Strains S0111_Mtb_Ec, S2190_Mtb_Ec, S2192_Mtb_Ec, and S2209_Mtb_Ec show discordant resistance results with KvarQ and PhyReSse (DST sensitive but genomic resistant characterised).

Diversity in genotypic resistance profile from the isolates was observed among the drug assessed, TB-Profiler detected a highest proportion of resistance to isoniazid with 63 isolates (71.6%) and ethambutol with 24 isolates (27.3%). Similarly SAM-TB detected 64 isolates (72.7%) for rifampicin while the lowest resistance proportions were mainly described by KvarQ.

It is noteworthy that all programs detected resistance to second-line drugs that were not detected by DST. This may be associated with the slower growth of some of the isolates, with these being at the limit of the resistance detection range. In particular, all programs detected 2 to 5 isolates had mutations associated with resistance to kanamycin, amikacin and capreomycin, none of which were previously detected by DST. As for fluoroquinolones, 10 to 13 isolates were characterised with sequences resistant to moxifloxacin and levofloxacin compared to 2 and 7 detected by DST, respectively.

Overall, KvarQ detected less resistance and PhyReSse, TB-Profiler and SAM-TB were the programs that detected more (Table 3).

**Table 3.** Comparison of the number of cases of resistance detection by phenotypic (DST) and genotypic web-based tools for ten TB anti-drugs

| Treatment drugs | DST<br>n (%) | Mykrobe<br>n (%) | TB-profiler<br>n (%) | PhyReSse<br>n (%) | KvarQ<br>n (%) | SAM-TB<br>n (%) |
| --- | --- | --- | --- | --- | --- | --- |
| Isoniazid | 63 (71.6) | 61 (69.3) | 63 (71.6) | 61 (69.3) | 59 (67.1) | 61 (69.3) |
| Rifampicin | 66 (75.0) | 63 (71.6) | 63 (71.6) | 64 (72.7) | 58 (65.9) | 64 (72.7) |
| Ethambutol | 11 (12.5) | 21 (23.9) | 24 (27.3) | 20 (22.7) | 13 (14.7) | 22 (25.0) |
| Pyrazinamide | 12 (13.6) | 14 (15.9) | 21 (23.9) | 14 (15.9) | 15 (17.0) | 21 (23.9) |
| Streptomycin | 16 (18.2) | 21 (23.9) | 32 (36.4) | 31 (35.2) | 26 (29.5) | 31 (35.2) |
| Kanamycin | 0 (0.0) | 3 (3.4) | 5 (5.7) | 3 (3.4) | 2 (2.3) | 5 (5.7) |
| Amikacin | 0 (0.0) | 3 (3.4) | 3 (3.4) | 3 (3.4) | 2 (2.3) | 3 (3.4) |
| Capreomycin | 0 (0.0) | 3 (3.4) | 3 (3.4) | 3 (3.4) | 2 (2.3) | 3 (3.4) |
| Levofloxacin | 7 (8.0) | 11 (12.5) | 13 (14.8) | 11 (12.5) | 10 (11.4) | 13 (14.8) |
| Moxifloxacin | 2 (2.3) | 11 (12.5) | 13 (14.8) | 11 (12.5) | 10 (11.4) | 13 (14.8) |

### Polymorphisms associated with drug resistance identified by whole genome sequencing and web-based tools

Through whole genome sequencing of the 88 isolates and using the five web tools mentioned above, we found a total of 59 mutations, of these, 5 were in promoter regions, 5 in intergenic regions and 49 in coding regions incurring changes in the reading frame. Most of them correspond to single nucleotide changes, but 8 insertions and 7 deletions were detected. Three of the deletions found corresponded to regions of considerable length, one of 193 nt in the *gid* gene and anothers of 891 and 833 nt affecting the *pncA* and *Rv2041c-Rv2042c* genes. The *pncA* and *rpoB* genes that are associated with resistance to pyrazinamide and rifampicin had the highest number of mutations. For *pncA*, 13 mutations and two insertions of 4 nt each and one deletion of 833 nt were detected. For *rpoB*, 13 nonsynonymous mutations were detected, several of which were present in the same codon. In fact position 445 of the *rpoB* gene is the most variable codon (generating 5 different configurations) followed by position 406 of the *embB* gene (with three different mutations). In addition, the *embB, katG, gyrA* and *rrs* genes, which conferred resistance to ethambutol, isoniazid, fluoroquinolones and streptomycin, respectively, had at least 4 mutations (Table 4, S3 Table).

**Table 4.**
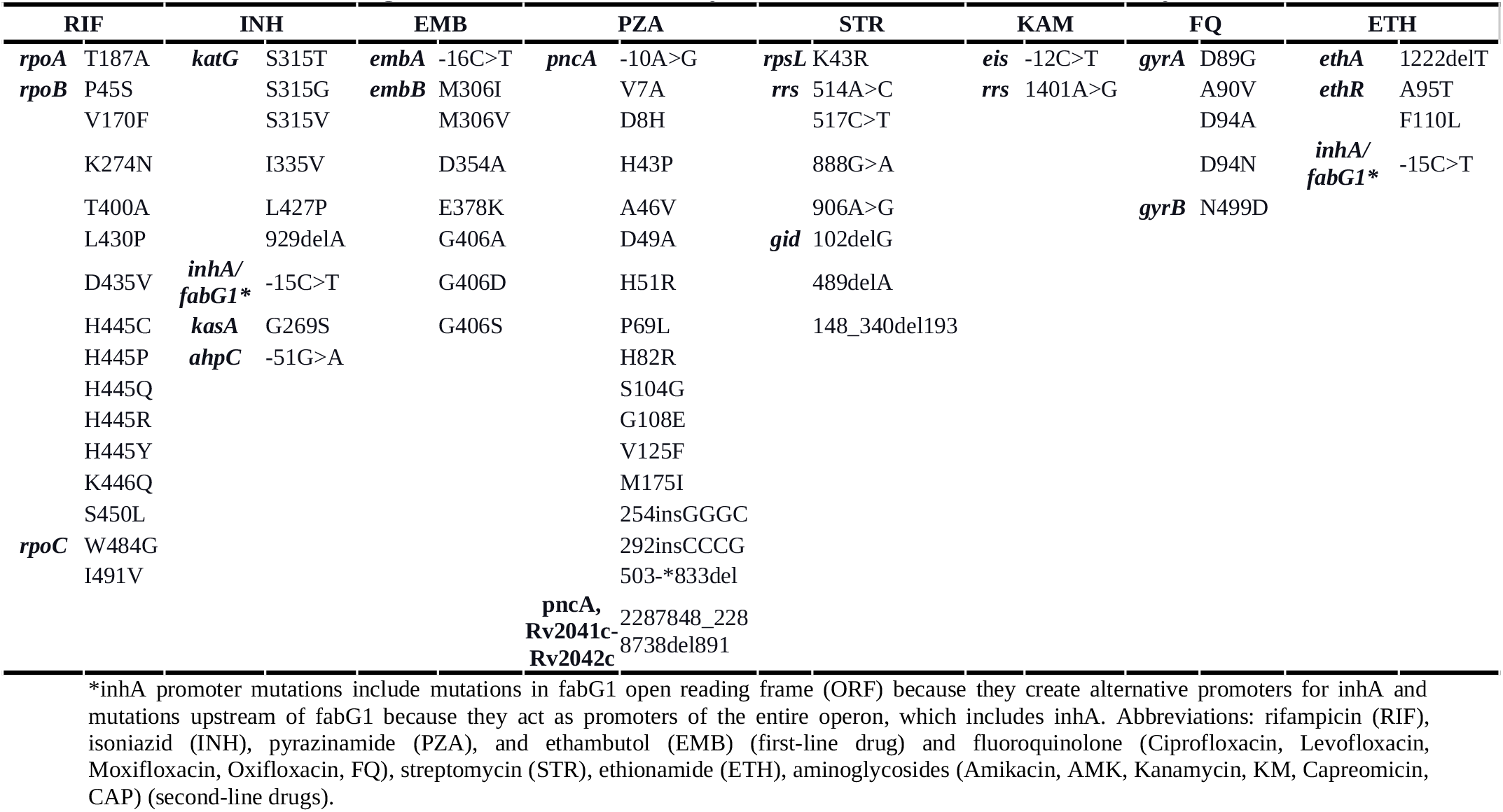
Mutational changes in resistant isolates of *Mycobacterium tuberculosis* identified by WGS

### Polymorphisms conferring Rifampicin resistance

From 66 isolates defined phenotypic resistant to rifampicin, mutations in *rpoB, rpoC* and *rpoA* genes were frequently identified by WGS. *rpoB* is a prime target for characterising resistance-related mutations because it has a hot-spot region, whereas *rpoA* and *rpoC* are genes with mutations considered compensatory mutations. In our data set, the S450L *rpoB* mutation was the most prevalent (39/64) followed by D435V (6/64), V170F (3/64), D435Y, H445C/R/Y, L430P, T400Ala (1/64 each). One isolate showing the V170F and L430P mutations and three isolates with the L430P and H445Q mutations were also identified. TB-profiler and SAM-TB were able to identify one isolate (S0555_MTb_Ec) with H445P and K446Q mutations, in the meantime SAM-TB was able to identify in one isolate (S0506_MTb_Ec) the mutation S450L plus K274N while the others only detected S450L. KvarQ was able to identify isolates with the compensatory mutations W484G and I491V in the *rpoC* and T187A mutation in the *rpoA* gene. Among sensitive isolates, none resistance mutation was characterised.

### Polymorphisms conferring Isoniazid and Ethionamide resistance

At least one known isoniazid-resistance mutation was found in 61 out of 63 clinical isolates where isoniazid resistance was reported from phenotypic susceptibility testing. For isolates S1135_Mtb_Ec and S1205_Mtb_Ec with resistance detected by DST, 426C>T mutation was found but none previously reported for isoniazid. On the contrary, the isolate S1136_Mtb_Ec reported sensitive by DST showed one resistant-related mutation. Mutation S315T in the katG gene was the most prevalent (43/63), followed by S315G (12/63, which was detected only by Mykrobe) and S315N detected only by TB-profiler, PhyReSse and KvarQ, in this case, Mykrobe detected this mutation how S315V.

Five isolates presented the mutation -15C>T in the gene fabG1/inhA promoter related to isoniazid/ethionamide resistance. TB-profiler, PhyReSse and SAM-TB were able to report other ethionamide resistance, in the *ethR* gene A95T and F110L mutations were identified for one isolate and in the *ethA* gene the 1222delT deletion was detected in three isolates.

For three isolates that presented resistance to isoniazid detected by DST, no canonical mutations related to this resistance were detected. Of these, the isolate S0889_Mtb_Ec presents mutations in the katG gene (L427P) and at the ahpC promoter (−51G>A). For the other two isolates, no known mutations related to isoniazid resistance were found, nor do they have common mutations. The mutation in the kasA gene (G269S) was present in three isolates reported as sensitive by DST and twenty-one as MDR.

### Polymorphisms conferring Ethambutol resistance

Ethambutol affects the *embCAB* gene locus which is involved in the biosynthesis of the cell wall components arabinogalactan and lipoarabinomannan. The resistant strains are principally characterised by reported canonical mutations at codons 306, 406 and 497 within the embB gene. In our study, we determined the mutations M306I/V, D354A, and G406A/D/S into the *embB* gene and -16C>T substitution in the *embA* gene. M306I was the most prevalent mutation (10/22) but also could be detected in combination with the mutations G406D and D354A followed by M306V and G406S (4/22 each). In particular, it is noteworthy that mutations in the embB and embA genes were found in 11 isolates characterised as sensitive to this drug by DST.

### Polymorphisms conferring Pirazinamide resistance

Pyrazinamide (PZA) is an important first-line anti-TB drug, as a prodrug, PZA is transformed by *pncA*-encoded pyrazinamidase into pyrazinoic acid. Resistance to pyrazinamide has been determined mainly by mutations in the *pncA* gene, although mutations in the *rpsA* and *panD* genes have also been identified in minor proportions. In our data, we identified different mutations in the *pncA* gene, including 8 insertions, 1 deletion, and 14 single nucleotide polymorphisms. These mutations were scattered throughout the entire length of the *pncA* gene, including the promoter region, being the more representative H51R, H82R, V125F (3/21) followed by G108E (2/21), and uniquely found mutations, V7A, D8H, H43P, A46V, D49A, P69L, S104G, L172P, M175I and the substitution -10A>G in the promoter region. One isolate has a large deletion of 891 nt spanning the pncA, Rv2041c-Rv2042c genes and another isolate has an 833 nt deletion within the pncA gene (503-*del833). Insertions 292GC, 293CC, 294CC and 295GG was observed in the isolate S0036_Mtb_Ec and 254TG, 255GG, 256GG and 257CC in the isolate S0040_Mtb_Ec. From eight isolates characterised as pyrazinamide-sensitive by DST were determined mutations in the gene *pncA*. Two isolates had mutations in the *rpsA* gene but were not related to resistance to pyrazinamide.

### Polymorphisms conferring Fluoroquinolones resistance

Fluoroquinolones (FQ), are critical to the treatment of MDR-TB, considered an important second-line anti-TB drug. Phenotypic resistance is associated with mutations in the quinolone resistance-determining region (QRDR) of DNA subunits A (*gyrA*) and B (*gyrB*). The most common mutations appear in codons 88, 90, 91, and 94 of the *gyrA* gene, and in codons 500 and 538 of the *gyrB* gene. By WGS from our analysis, we determined 13 isolates with mutations associated with resistance to FQ, being the more frequent A90V (8/13) followed by D94N (3/13) and D89G (1/13) all in the *gyrA* gene, one isolate had a combination of two mutations, D94A in the gyrA gene plus N499D in gyrB.. In one isolate identified as resistant to Levofloxacin plus Moxifloxacin by DST, no resistance-related mutation could be determined by WGS. No mutations related to this resistance were detected in any isolate identified as sensitive by DST.

### Polymorphisms conferring Aminoglycoside resistance

Aminoglycosides have played a prominent role in the treatment of tuberculosis (TB) from the first trial to the present, being streptomycin, kanamycin, amikacin, and capreomycin used under different considerations (28). Polymorphisms in the *rpsL* gene (ribosomal protein S12) were associated with high streptomycin resistance levels, while those in the *rrs* (16S ribosomal RNA) or the *gid* genes were linked to intermediate to high and low STR resistance levels, respectively. Of 28 isolates characterised as streptomycin resistant by WGS, 16 had the K43R mutation in the rpsL gene and only one had the K88R mutation in the same gene, being the least frequent. The substitutions 492C>T (8/28), 514A>C (3/28), 517C>T, 888G>A and 906A>G were reported in the *rrs* gene. Furthermore the deletion 102delG, 489delA and 148_340del193 were presented in the *gid* gene. Notably, four isolates characterised as streptomycin-sensitive by DST presented the known substitution 492C>T in the *rrs* gene.

The second-line injectable drugs kanamycin, amikacin, and capreomycin are crucial to the prevention of XDR-TB and effective in the MDR-TB treatment, for this the appropriate use is recommended (29). Single nucleotide polymorphism and variations in the *rrs* and *eis* genes are the most frequent in resistant strains. The classical mutations to confer resistance are 1401A>G, 1402C>T, and 1484G>T in the *rrs* gene and -37G>T, -14C>T, -12C>T, and -10G>A in the *eis* gene. Through DST our isolates were characterised as sensitive to kanamycin, amikacin, and capreomycin, however by WGS we identified three isolates that presented the classical 1401A>G in the *rrs* gene and two isolates with the -12C>T mutation in the *eis* gene.

## Discussion

Although sequencing-based diagnostic information for *M. tuberculosis* has been available for some years and methodological advances have made possible the use of direct whole-genome sequencing of *M. tuberculosis* for the accurate prediction of drug resistance, this study assesses and compares the use of whole genome sequencing and user-friendly web tools to infer resistance in tuberculosis drug susceptibility surveillance in Ecuador. The isolates came from 9 cities with Guayaquil being the most representative city (59.1%); the high number of tuberculosis cases detected in this city is possibly due to the high mobility from other provinces for trade or work, in addition to the location of the leading health centres for the monitoring of this pathogen.

Providing adequate treatment based on rapid detection of resistant strains, as well as the identification of transmission clusters are important elements that would lead to a decrease in tuberculosis (4), for which the use of procedures that reduce the time of diagnosis is recommended. Whole-genome sequencing (WGS) is a powerful method for detecting drug resistance, genetic diversity, and transmission dynamics of *M. tuberculosis*. Moreover, in recent times, the World Health Organisation (WHO) publishes updates of guidelines for TB treatment including drug-susceptible, single drug-resistant, multidrug-resistant (MDR-TB) and extensively drug-resistant TB (XDR-TB)(23) which has allowed to improve the pipelines within the web-based tools. Despite the improvements in the WGS protocols for TB diagnosis that simultaneously allows, species differentiation, detection of drug resistance-associated mutations, and phylogenetic/clustering analyses to guide contact investigations (30–32), analysis of genomic sequencing data remains an obstacle to the routine use of WGS technology in clinical tuberculosis. This is mainly because it requires bioinformatics expertise, high-performance computing, funding, and training that are not readily available in most clinical laboratories (24,33).. In our analysis, 97% of phenotypic resistant isolates were correctly detected by whole genome sequencing and 18.2% of phenotypic sensitive isolates reported mutations associated with resistance. The detection of mutation in DST-negative isolates was specifically due to the presence of the substitution 492C>T in the *rrs* gene associated with streptomycin resistance. The reason for discordant results could be due to critical concentration in some DST systems, mutation outside the DR regions, silent mutations and heteroresistance (34,35).

WGS has previously been evaluated as a useful tool for quantifying transmission clusters, within which a threshold of 12 SNPs maximises agreement between epidemiological research and genomic data (36,37). In our study, using 12 SNPs we identified sixteen mostly medium-sized transmission clusters (3 - 5 isolates) indicating local aggregations. Furthermore, we showed a relatively high transmission rate (54/88, 61.4%). These results differ from previous local studies where a low transmission rate was reported when using MIRU-VNTR (38,39) or using WGS for the localities of Valencia (24%) (40) and Los Angeles (25%) (41). This could be due to a large percentage of cluster of two isolates (43.8%) in our results or the highest SNPs umbral used in other studies (15 SNPs).

Over the last few years some TB-specific genome browsers and WGS analysis tools such as TBProfiler, KvarQ, TGS-TB, Mykrobe Predictor TB, CASTB, PhyTB, ReSeqTB-UVP, GenTB, PhyResSE, SAM-TB have been developed to genotyping and drug resistance identification. However, despite their usefulness have not been widely used within surveillance programs in many countries, including Ecuador. By evaluating the accuracy, sensitivity and specificity of five tools for inferring resistance of M. tuberculosis isolates compared to phenotypic tests, we show that globally PhyReSse, Mykrobe and SAM-TB have better ability to identify resistance in *M tuberculosis* isolates (97.0% sensitivity, 100.0% specificity, and 97.7% accuracy). In particular SAM-TB, the most recently developed tool, includes features of other two pipelines and improves the detection and interpretation of resistance.

In *M tuberculosis* drug resistance is mediated through mutations in specific gene targets. Therefore, the key is to identify those SNPs which are responsible for or strongly associated with resistance (8,25,42,43). Previous studies have indicated canonical mutations associated with resistance (32,44–50) as well as deletions that affect gene functionality (51) or promote bacterial transmission (52). In our study, only 19 resistance-associated genes were identified among 42 candidate genes, involving a total of 73 mutations. The rate of dominant mutations among the drugs evaluated was between 14% and 68%, specifically RIF (60.9%), INH (68.3%), EMB (45.5%), PZA (14.3% each), FQ (61.5%), STR (57.1%). In addition, interesting large deletions related to the pncA and gid genes were detected.In turn, non-resistance-conferring mutations were detected in the katG (R463L) and gyrA (E21Q, S95T, G668D) genes that may be useful for the evolutionary characterization of the *M. tuberculosis* genome. (8,53).

The resistance to rifampicin generally is exhibited by mutations in the 81-bp rifampicin resistance-determining region (RRDR) of the *rpoB* gene, also known as hotspot region, have been accurate predictors of rifampicin resistance in many studies (45,54,55). In fact, 60.9% of the RIF-resistant isolates possessed the S450L mutation in rpoB, in agreement with previous studies that reported a higher prevalence of this mutation associated with high levels of resistance, as well as H445Y and H445D (68,69). Other mutations that tend to cause moderate or low resistance (D435V, D435Y, H445L and L452P) were also detected, but at low proportions. In addition, we identified compensatory mutations that are associated with MDR transmission, such as W848G and I491V in the rpoC gene and T187A in the rpoA gene. (3,58–61).

Mutations in *katG, kasA, inhA, fabG1* genes and the *oxyR-ahpC* intergenic region are associated with conferring resistance to isoniazid (44,62,63), thus ethA/R mutations are associated with ethionamide resistance (48,64,65). Our results were in concordance with previous studies (44,63,66) due to the higher prevalence of mutations in *katG* (88.8%). Furthermore, it has been shown that mutations in the *ahpC* promoter conferred resistance by compensatory mechanism (67,68) like L427P in the *katG* gene (69,70). Resistance to ethambutol is due to mutations in the embCAB gene operon, which is involved in the biosynthesis of the cell wall components arabinogalactan and lipoarabinomannan (46,71–74). In agreement with previous studies, we showed that 95.5% of ethambutol resistance is due to SNPs in embB (M306I/V, D354A, G406A/D/S), and 9.1% to the -16C>T substitution in the embA gene. Similarly, all the mutations we found related to pyrazinamide resistance (95.2% SNP and 26.6% Indels in pncA) coincided with studies pointing to the importance of the promoter and the pncA gene (4,46,51,75,76).

Our results when applying WGS show that 92.3% of mutations were presented in the *gyrA* gene, especially in the quinolone resistance-determining region (QRDR) D89G, A90V and D94N codons and one isolate presented combined mutations D94A into *gyrA* gene plus N499D in *gyrB*. This is in concordance with various studies that reported mutations in the conserved QRDR region of gyrase genes (gyrA and gyrB) which changes the structure of the drug binding pocket (QBP) and results in cross-resistance to all FQs (64,77–79). One resistant isolate did not have gyrA or gyrB mutations. This may be explained by the fact that additional mechanisms such as efflux pumps may be involved in generating resistance fluoroquinolones (80).

A high proportion of STR resistance was observed in this study, which 60.7% of mutations was in the *rpsL* gene, agreeing that the k43R mutation is the most recurrent streptomycin resistance-associated mutation (32,79,81). In the *rrs* gene, the substitutions identified at positions 492, 514, 517, 888 and 906 previously have been associated with streptomycin resistance in TB (8,62,82,83); even though the 1401A>G, 1402C>T, and 1484G>T have been associated with cross-resistance to kanamycin, capreomycin or amikacin (84–86), only 1401A>G mutation was presented in 7.1% of isolates. Variations in the promoter (−37G>T, -14C>T, -12C>T, and -10G>A) of the *eis* gene have been associated with a low resistance to kanamycin (87,88). Our results support this finding as two isolates with the -12C>T mutation in the *eis* promoter.

This is the first extensive overview of the resistance of *M. tuberculosis* in Ecuador, applying whole genome sequencing and web-based bioinformatic tools, showing the ability of molecular assays to detect drug resistance in tuberculosis strains that allow adequate monitoring to generate health policies. They should therefore be applied more routinely to improve TB strain surveillance programs in South American countries widely affected by TB.

## Material and methods

Eighty-eight clinical isolates were provided by private laboratories and the National Reference of Micobacterias of the National Institute of Public Health Research “Leopoldo Izquieta Pérez” (INSPI-LIP for spanish acronyms) in Guayaquil between 2019 to 2021. These isolates were previously identified as multidrug-resistant by DST agar or Bactec MGIT 960 System protocol (28). The resistance pattern of first-line and second-line anti-tuberculosis drugs were performed according to the proportional method by Bactec MGIT 960 System protocol in 67 samples with critical concentrations of drugs as follow: rifampicin, 1.0 μg/mL; isoniazid, 0.1 μg/mL; ethambutol, 5.0 μg/mL; kanamycin, 2.5 μg/mL, capreomycin, 1.0 μg/mL, amikacin, 1.0 μg/mL, 1.0 µg/mL for Levofloxacin and 1.0 µg/mL for Moxifloxacin, however in 21 sample the resistance was determined by DST agar (29), the critical concentrations of anti-TB drugs for the DST assays were as follows: rifampicin, 40.0 μg/mL; isoniazid, 0.2 μg/mL; ethambutol, 0.4 μg/mL; streptomycin, 4.0 μg/mL, and 200 µg/mL for pyrazinamide (pyrazinamidase assay); from results of all isolate the resistance profile was defined according to WHO recommendations.

### DNA extraction and sequencing

Genomic DNA was extracted from 88 isolates of *Mycobacterium tuberculosis* grown in Lowenstein-Jensen medium by two different protocols. In 21 isolates the DNA was extracted by the CTAB method (30,31) while for others 67 were obtained by the PureLink DNA Mini Kit (Fisher Scientific, Pennsylvania, USA) according to manufacturer instructions. Each extracted DNA was quantified by a Qubit 4.0 fluorometer (Invitrogen, Carlsbad, CA, USA.). DNA samples that fulfil the quality standards in terms of integrity, purity and quantity were sequenced. Genomic DNA libraries were prepared for whole genome sequencing using the Tagmentation-based library prep kit according to the manufacturer’s instructions (Illumina Inc, San Diego, CA, USA). Next, libraries with different indices were multiplexed and loaded on an Illumina MiniSeq platform with High Output Reagent Kit, according to the manufacturer’s instructions (Illumina, San Diego, CA, USA). Sequencing was carried out using a 2 × 150 paired-end (PE) configuration.

### Bioinformatic analyses

Using the Galaxy platform (https://usegalaxy.org/) reads were classified by Kraken version 2 (32) to detect possible contamination or the presence of other mycobacteria. In addition, FastQC version 0.11.9 (33) and Trimmomatic version 0.38 (34) were used to control the quality and trim the low-quality ends of the reads, respectively. In particular, a sliding window was used to trim sequences with an average quality value lower than 20. The high-quality reads were mapped to *M. tuberculosis* strain HR37v (NC_000962.3) using the BWA-MEM (35). A very strict criterion was used to discard isolates with sequences that were not fully reliable. Thus, isolates with sequencing depth of less than 20X or with a reference coverage of less than 90% were not considered for the following analyses.

For phylogenetic analysis, clean reads were used as input in MTBseq pipeline (97) to obtain i) the sub-lineage classification, ii) the transmission groups identification and iii) the SNPs-based alignment for phylogenomic analysis. *Mycobacterium microti* Maus III (Accession number: ERR4618952) was used as an outgroup to obtain the rooted tree. Substitution model was calculated with ModelTest-NG v0.1.7 (98). Phylogenetic reconstruction was performed with RAxML-NG (99) using the maximum likelihood method and a bootstrap cutoff = 0.01. Visualisation of the phylogenetic tree was achieved with iTOL v6.6 (100). The transmission groups were determined by evaluating a distance of 12 SNPs between strains. The transmission cluster sizes were classified according to the number of shared isolates as follows: small if it had less than three isolates, medium if it composed between three to five isolates and large if it had more than five isolates.

### Predicting susceptibility and drug resistance

Web-based tools such as TB-Profiler v2.8.13 (https://tbdr.lshtm.ac.uk/), PhyReSse v1.0 (The <u>Phy</u>lo-<u>Res</u>istance-<u>S</u>earch-<u>E</u>ngine) (https://bioinf.fz-borstel.de/mchips/phyresse/), Mykrobe v0.10.0 (https://www.mykrobe.com/), KvarQ v0.12 (https://gap.tbportals.niaid.nih.gov/#/dashboard/home) and SAM-TB (https://samtb.szmbzx.com/index), were used for predicted the canonical mutation in genes related to resistance such as rifampicin (RIF), isoniazid (INH), pyrazinamide (PZA), and ethambutol (EMB) (first-line drug) and fluoroquinolone (Ciprofloxacin, Levofloxacin, Moxifloxacin, Oxifloxacin, FQ), streptomycin (STR), ethionamide (ETH), aminoglycosides (Amikacin, AMK, Kanamycin, KM, Capreomicin, CAP) (second-line drugs). All programs were run under the default parameters using the high-quality reads from the 88 samples.

### Statistical analysis

Sensitivities, specificities, positive predictive values (PPV) and negative predictive values (NPV) global and drug dependent were calculated using the statistical software MedCalc® Statistical Software version 20.210 (MedCalc Software Ltd, Ostend, Belgium; https://www.medcalc.org; 2022) with a 95% confidence interval using as a reference (gold standard) the diagnostic results based on DST information.

## Ethics statements

The study was approved by the ethics committee of University Espíritu Santo (code 2022-001A) certified by the Ministry of Public Health from Ecuador following guidelines from the Declaration of Helsinki. All samples were anonymized, and no data on the patients were made available. The present work is based on the INSPI-LIP permission to use the positive samples.

## Data availability

Raw sequence reads of all *Mycobacterium tuberculosis* isolates subjected these WGS analysis were deposited in the Sequence Read Archive (SRA) (http://www.ncbi.nlm.nih.gov/sra) accessible under the BioProject number PRJNA827129.

## Acknowledgements

We acknowledge the FCI Research program of the University of Guayaquil for the partial financial support of the procurement of the reagents to the MiniSeq sequencing machine that were used to run the samples. We thank the private laboratories and National Reference of Micobacterias of INSPI-LIP for supporting the isolate analysed in this study. We also thank the Omics Lab of Universidad Espiritu Santo for permitting the use of the MiniSeq sequencing machine employed to run the samples.We also thank the Bioinformatics Unit of the Institute Pasteur de Montevideo for their assistance and support.

## Authors’ contributions

GML conceived and designed the study, performed the collection of isolates, sample processing, data analysis (bioinformatic processing of the raw sequencing data) and drafted the manuscript, JCPG performed the collection of isolates on the National Reference of Micobacterias, PMMP, DAM, KMM, JCFC and JCGP performed sample processing (library prep and sequencing), data analysis and review the draft manuscript, EGM developed statistical analysis, CLC provided advice on data analysis, LB conceived and designed the study, wrote the main manuscript and review it; The funding acquisition was performed by GM and DAM. All authors read and approved the final manuscript.

